# A genome-wide screen for genes affecting spontaneous direct-repeat recombination in *Saccharomyces cerevisiae*

**DOI:** 10.1101/2020.02.11.943795

**Authors:** Daniele Novarina, Ridhdhi Desai, Jessica A. Vaisica, Jiongwen Ou, Mohammed Bellaoui, Grant W. Brown, Michael Chang

## Abstract

Homologous recombination is an important mechanism for genome integrity maintenance, and several homologous recombination genes are mutated in various cancers and cancer-prone syndromes. However, since in some cases homologous recombination can lead to mutagenic outcomes, this pathway must be tightly regulated, and mitotic hyper-recombination is a hallmark of genomic instability. We performed two screens in *Saccharomyces cerevisiae* for genes that, when deleted, cause hyper-recombination between direct repeats. One was performed with the classical patch and replica-plating method. The other was performed with a high-throughput replica-pinning technique that was designed to detect low-frequency events. This approach allowed us to validate the high-throughput replica-pinning methodology independently of the replicative aging context in which it was developed. Furthermore, by combining the two approaches, we were able to identify and validate 35 genes whose deletion causes elevated spontaneous direct-repeat recombination. Among these are mismatch repair genes, the Sgs1-Top3-Rmi1 complex, the RNase H2 complex, genes involved in the oxidative stress response, and a number of other DNA replication, repair and recombination genes. Since several of our hits are evolutionary conserved, and repeated elements constitute a significant fraction of mammalian genomes, our work might be relevant for understanding genome integrity maintenance in humans.

## INTRODUCTION

Homologous recombination (HR) is an evolutionarily conserved pathway that can repair DNA lesions, including double-strand DNA breaks (DSBs), single-strand DNA (ssDNA) gaps, collapsed replication forks, and interstrand crosslinks, by using a homologous sequence as the repair template. HR is essential for the maintenance of genome integrity, and several HR genes are mutated in human diseases, especially cancers and cancer-prone syndromes (Krejci et al., 2012; Symington et al., 2014). HR is also required for meiosis (Hunter, 2015) and is important for proper telomere function (Claussin and Chang, 2015). The yeast *Saccharomyces cerevisiae* has been a key model organism for determining the mechanisms of eukaryotic recombination. Our current understanding of the HR molecular pathway comes mainly from the study of DSB repair. However, most mitotic HR events are likely not due to the repair of DSBs (Claussin et al., 2017), and can be triggered by diverse DNA structures and lesions, including DNA nicks, ssDNA gaps, arrested or collapsed replication forks, RNA-DNA hybrids and noncanonical secondary structures (Symington et al., 2014). An essential intermediate in recombination is ssDNA, which, in the case of a DSB, is generated by resection of the DSB ends by nucleases. Rad52 stimulates the loading of Rad51 onto ssDNA, which in turn mediates homologous pairing and strand invasion, with the help of Rad54, Rad55, and Rad57. After copying the homologous template, recombination intermediates are resolved with the help of nucleases and helicases, and the HR machinery is disassembled (Symington et al., 2014).

While HR is important for genome integrity, excessive or unregulated recombination in mitotic cells can be deleterious. Indeed, even though HR is generally considered an error-free DNA repair pathway, outcomes of HR can be mutagenic. For instance, single strand annealing (SSA) occurring between direct repeats results in the deletion of the intervening sequence (Bhargava et al., 2016), while recombination between ectopic homolog sequences can lead to gross chromosomal rearrangements (Heyer, 2015). Mutations and chromosomal aberrations can be the outcome of recombination between slightly divergent DNA sequences, a process termed “homeologous recombination” (Spies and Fishel, 2015). Allelic recombination between homologous chromosomes can lead to loss of heterozygosity (LOH) (Aguilera and García-Muse, 2013). Finally, the copying of the homologous template occurs at lower fidelity than is typical for replicative DNA polymerases, resulting in mutagenesis (McVey et al., 2016). For these reasons, the HR process must be tightly controlled, and spontaneous hyper-recombination in mitotic cells is a hallmark of genomic instability (Aguilera and García-Muse, 2013; Heyer, 2015).

Pioneering mutagenesis-based screens led to the identification of hyper-recombination mutants (Aguilera and Klein, 1988; Keil and McWilliams, 1993). Subsequently, several systematic screens were performed with the yeast knockout (YKO) collection to identify genes whose deletion results in a spontaneous hyper-recombinant phenotype. In particular, Alvaro et al. screened an indirect phenotype, namely elevated spontaneous Rad52 focus formation in diploid cells, which led to the identification of hyper-recombinant as well as recombination-defective mutants (Alvaro et al., 2007). A second screen for elevated Rad52 foci in haploid cells identified additional candidate recombination genes (Styles et al., 2016), although the recombination rates of these were not assessed directly. A distinct screen of the YKO collection measured elevated spontaneous LOH events in diploid cells, which arise through recombination between homologous chromosomes or as a consequence of chromosome loss (Andersen et al., 2008). Here we describe two systematic genome-scale screens measuring spontaneous recombination in haploid cells, since the sister chromatid is generally a preferred template for mitotic recombination relative to the homologous chromosome, both in yeast and mammalian cells (Johnson and Jasin, 2000; Kadyk and Hartwell, 1992). We use a direct-repeat recombination assay (Smith and Rothstein, 1999), because recombination between direct repeats can have a significant impact on the stability of mammalian genomes, where tandem and interspersed repeated elements, such as LINEs and SINEs, are very abundant (George and Alani, 2012; López-Flores and Garrido-Ramos, 2012).

Recombination rate screens were performed both with the classical patch and replica-plating method and with our recently developed high-throughput replica-pinning technique, which was designed for high-throughput screens involving low-frequency events (Novarina et al., 2020). High-throughput replica-pinning is based on the concept that, by robotically pinning an array of yeast strains many times in parallel, several independent colonies per strain can be analysed at the same time, giving a semi-quantitative estimate of the rate at which a specific low-frequency event occurs in each strain. We used both approaches to screen the YKO collection with the direct-repeat recombination assay. Bioinformatic analysis and direct comparison of the two screens confirmed the effectiveness of the high-throughput replica-pinning methodology. Together, we identified and validated 35 genes whose deletion results in elevated spontaneous direct-repeat recombination, many of which have homologs or functional counterparts in humans.

## MATERIALS AND METHODS

### Yeast strains and growth conditions

Standard yeast media and growth conditions were used (Sherman, 2002; Treco and Lundblad, 2001). All yeast strains used in this study are derivatives of the BY4741 genetic background (Brachmann et al., 1998) and are listed in Supporting Information, Table S1.

### Patch and replica-plating screen

To create a recombination assay strain compatible with Synthetic Genetic Array (SGA) methodology (Kuzmin et al., 2016), the *leu2ΔEcoRI-URA3-leu2ΔBstEII* direct repeat recombination reporter (Smith and Rothstein, 1999) was introduced into Y5518 by PCR of the *LEU2* locus from W1479-11C, followed by transformation of Y5518 and selection on SD-ura. Correct integration was confirmed by PCR, and the resulting strain was designated JOY90. JOY90 was then crossed to the *MAT***a** yeast knockout (YKO) collection ((Giaever et al., 2002); gift of C. Boone, University of Toronto), using SGA methodology (Kuzmin et al., 2016). Following selection on SD-his-arginine-lysine-uracil+G418+ClonNat+canavanine+thialysine, the resulting strains have the genotype *MAT***a** *xxxΔ::kanMX mfa1Δ::MFA1pr-HIS3 leu2ΔEcoRI::URA3-HOcs::leu2ΔBstEII his3Δ1 ura3Δ0 met15Δ0 lyp1Δ can1Δ::natMX*, where *xxxΔ::kanMX* indicates the YKO gene deletion in each resulting strain.

Each YKO strain carrying the recombination reporter was streaked for single colonies on SD-ura. Single colonies were then streaked in a 1 cm x 1 cm patch on YPD, incubated at 30°C for 24 h, and then replica-plated to SD-leu to detect recombination events as papillae on the patch. RDY9 (wild-type) and RDY13 (*elg1Δ::kanMX*; positive control) were included on each plate. The papillae on SD-leu were scored by visual inspection relative to the control strains, yielding 195 positives (Table S2). The 195 positives were tested in a fluctuation test of 5 independent cultures, and those with a recombination rate of at least 2×10^-5^ (approximately twofold greater than that of RDY9) were identified (43 strains; Table S2). Positives from the first fluctuation tests (except *slm3*Δ and *pex13*Δ, where rates could not be determined due to the large numbers of ‘jackpot’ cultures where all colonies had a recombination event) were assayed further, again with 5 cultures per fluctuation test. Thirty-three gene deletion mutants displayed a statistically supported increase in recombination rate (Table S2, Figure 1D), using a one-sided Student’s t-test with a cutoff of p=0.05.

**Figure 1.**
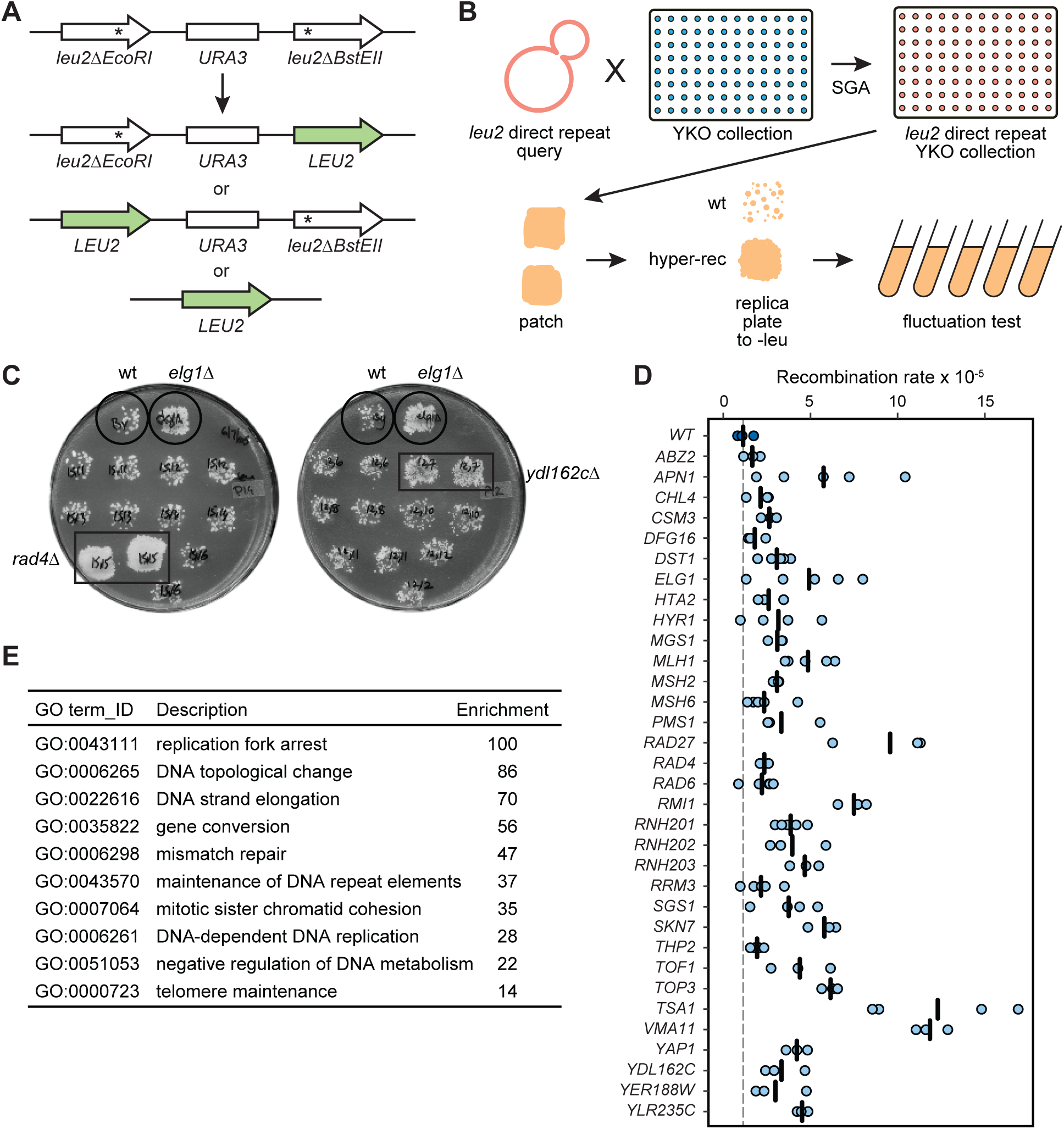
A genome-wide patching and replica plating screen for mutants with increased direct-repeat recombination. (**A**) The *leu2* direct-repeat recombination assay. Spontaneous recombination between two *leu2* heteroalleles, either through gene conversion or intra-chromosomal single strand annealing (SSA), yields a functional *LEU2* gene. (**B**) Schematic representation of the screen based on patching and replica plating. The *leu2* direct-repeat recombination cassette was introduced into the yeast deletion collection (YKO) by crossing the collection with a query strain containing the cassette. Haploid strains containing each gene deletion and the recombination cassette were isolated using SGA methodology. Each strain was patched on rich medium and replica-plated to selective medium, where hyper-recombinant mutants form papillae on the surface of the patch. Recombination rates were measured for positives from the patch assay using fluctuation tests. (**C**) Example plates from the patch assay. Each plate bears a negative control (wild type) and a positive control (*elg1Δ*). Two positive hits from the screen (*rad4Δ*, *ydl162cΔ*) are shown. (**D**) Recombination rates are plotted for the validated positives from the patch screen, alongside the wild-type strain. Each data point is from an independent fluctuation test, with n≥3 for each strain. The vertical bars indicate the mean recombination rate for each strain. (**E**) The top 10 statistically supported GO terms enriched in the hits from the patch assay screen are shown, with the -fold enrichment for each term.

### Fluctuation tests of spontaneous recombination rates

Fluctuation tests as designed by Luria and Delbrück (Luria and Delbrück, 1943) were performed by transferring entire single colonies from YPD plates to 4 ml of YPD liquid medium. Cultures were grown at 30°C to saturation. 100 µl of a 10^5^-fold dilution were plated on a fully supplemented SD plate and 200 µl of a 10^2^-fold dilution were plated on an SD-leu plate. Colonies were counted after incubation at 30°C for 3 days. The number of recombinant (leu+) colonies per 10^7^ viable cells was calculated, and the median value was used to determine the recombination rate by the method of the median (Lea and Coulson, 1949).

### High-throughput replica pinning screen

High-throughput manipulation of high-density yeast arrays was performed with the RoToR-HDA pinning robot (Singer Instruments). The *MAT***a** yeast deletion collection (EUROSCARF) was arrayed in 1536 format (each strain in quadruplicate). The *leu2ΔEcoRI-URA3-leu2ΔBstEII* marker to measure direct-repeat recombination (Smith and Rothstein, 1999) was introduced into the deletion collection through synthetic genetic array (SGA) methodology (Kuzmin et al., 2016) using the JOY90 query strain. The procedure was performed twice in parallel to generate two sets of the yeast deletion collection containing the *leu2* direct-repeat recombination reporter. Each plate of each set was then pinned onto six YPD+G418 plates (48 replicates per strain in total), incubated for one day at 30° and then scanned with a flatbed scanner. Subsequently, each plate was pinned onto SD-leu solid medium and incubated for two days at 30° to select recombination events. Finally, all plates were re-pinned on SD-leu solid medium and incubated for one day at 30° before scanning. Colony area measurement was performed using the ImageJ software package (Schneider et al., 2012) and the ScreenMill Colony Measurement Engine plugin (Dittmar et al., 2010), to assess colony circularity and size in pixels. Colony data was filtered to exclude artifacts by requiring a colony circularity score greater than 0.8. Colonies with a pixel area greater than 50% of the mean pixel area were scored for strains pinned to YPD+G418.

Following replica-pinning to SD-leu, colonies were scored if the pixel area was greater than 10% of the mean pixel area for the same strain on YPD+G418. For each deletion strain, the ratio of recombinants (colonies on SD-leu) to total colonies (colonies on YPD+G418) is the recombination frequency (Table S3). Strains where fewer than 10 colonies grew on YPD+G418 were removed from consideration, as were the 73 YKO collection strains carrying an additional *msh3* mutation (Lehner et al., 2007). The final filtered data is presented in Table S4.

### Gene Ontology enrichment analysis and functional annotation

GO term analysis was performed using the GO term finder tool (http://go.princeton.edu/) using a P-value cutoff of 0.01 and applying Bonferroni correction, querying biological process enrichment for each gene set. GO term enrichment results were further processed with REViGO (Supek et al., 2011) using the “Medium (0.7)” term similarity filter and simRel score as the semantic similarity measure. Terms with a frequency greater than 15% in the REViGO output were eliminated as too general. Gene lists used for the GO enrichment analyses are in Table 1, and the lists of enriched GO terms obtained are provided in Table S6. Human orthologues in Table 3 were identified using YeastMine (https://yeastmine.yeastgenome.org/yeastmine; accessed June 25, 2019). Protein-protein interactions were identified using GeneMania (https://genemania.org/; (Warde-Farley et al., 2010)), inputting the 35 validated hyper-rec genes, and selecting only physical interactions, zero resultant genes, and equal weighting by network. Network edges were reduced to a single width and nodes were annotated manually using gene ontology from the *Saccharomyces* Genome Database (https://www.yeastgenome.org). Network annotations were made with the Python implementation of Spatial Analysis of Functional Enrichment (SAFE) (Baryshnikova, 2016); https://github.com/baryshnikova-lab/safepy). The yeast genetic interaction similarity network and its functional domain annotations were obtained from (Costanzo et al., 2016). The genetic interaction scores for *YER188W*, *DFG16*, *VMA11*, and *ABZ2* were downloaded from the Cell Map (http://thecellmap.org/; accessed January 9, 2020),

**Table 1.**
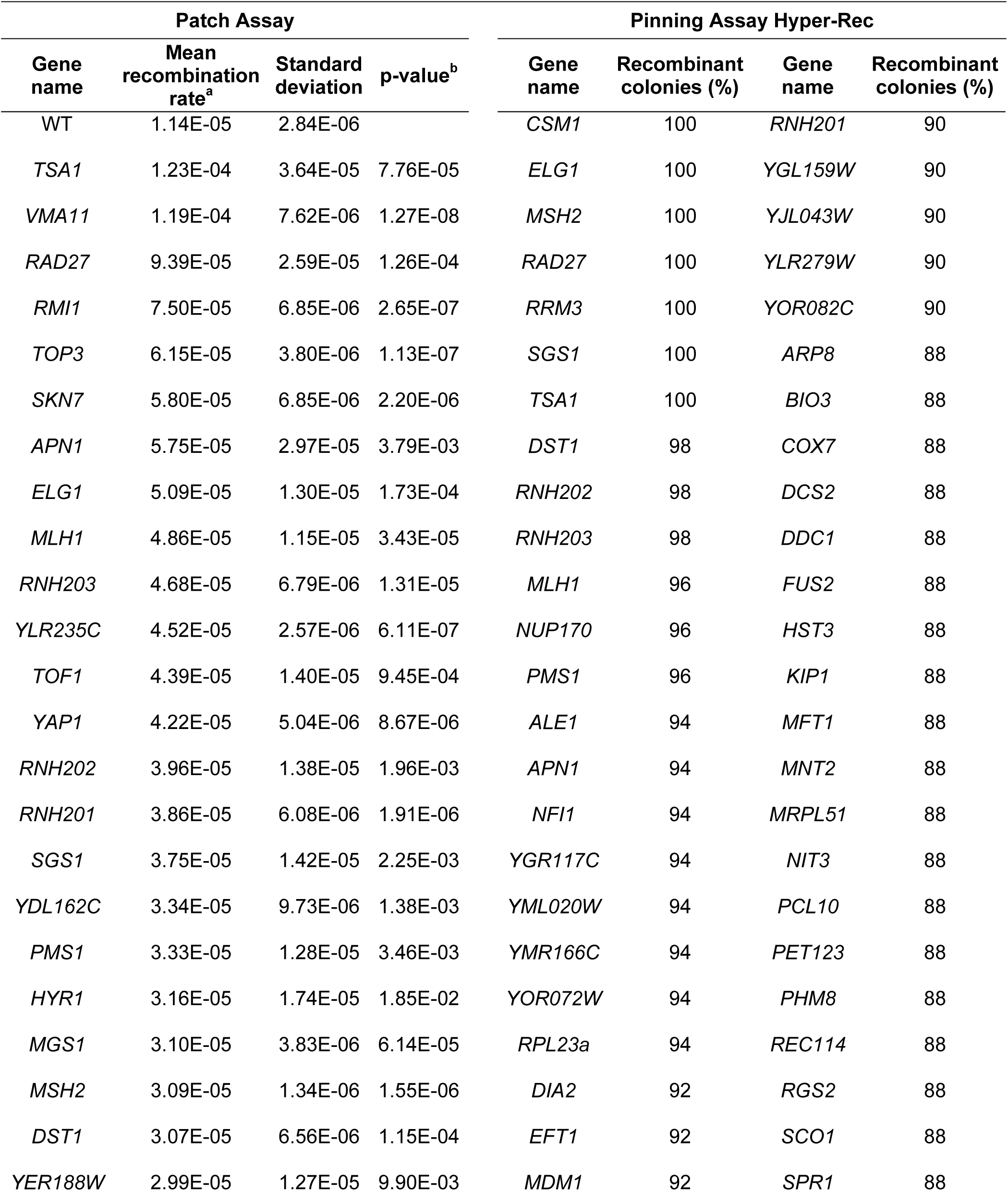
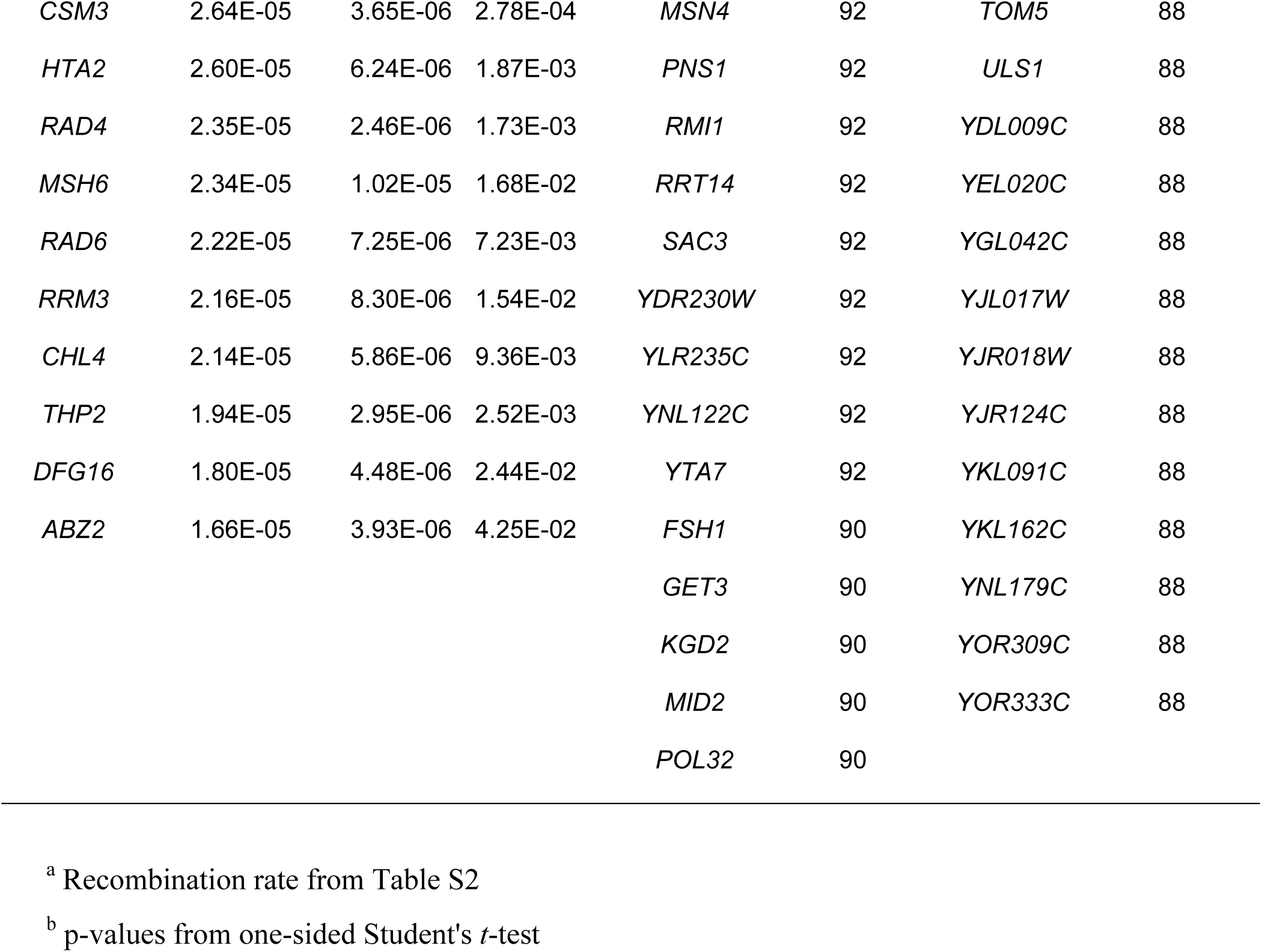
Hyper-recombination genes from the patch assay and pinning assay screens.

| Patch Assay |  |  |  | Pinning Assay Hyper-Rec |  |  |  |
| --- | --- | --- | --- | --- | --- | --- | --- |
| Gene name | Mean recombination rate<sup>a</sup> | Standard deviation | p-value<sup>b</sup> | Gene name | Recombinant colonies (%) | Gene name | Recombinant colonies (%) |
| WT | 1.14E-05 | 2.84E-06 |  | *CSM1* | 100 | *RNH201* | 90 |
| *TSA1* | 1.23E-04 | 3.64E-05 | 7.76E-05 | *ELG1* | 100 | *YGL159W* | 90 |
| *VMA11* | 1.19E-04 | 7.62E-06 | 1.27E-08 | *MSH2* | 100 | *YJL043W* | 90 |
| *RAD27* | 9.39E-05 | 2.59E-05 | 1.26E-04 | *RAD27* | 100 | *YLR279W* | 90 |
| *RMI1* | 7.50E-05 | 6.85E-06 | 2.65E-07 | *RRM3* | 100 | *YOR082C* | 90 |
| *TOP3* | 6.15E-05 | 3.80E-06 | 1.13E-07 | *SGS1* | 100 | *ARP8* | 88 |
| *SKN7* | 5.80E-05 | 6.85E-06 | 2.20E-06 | *TSA1* | 100 | *BIO3* | 88 |
| *APN1* | 5.75E-05 | 2.97E-05 | 3.79E-03 | *DST1* | 98 | *COX7* | 88 |
| *ELG1* | 5.09E-05 | 1.30E-05 | 1.73E-04 | *RNH202* | 98 | *DCS2* | 88 |
| *MLH1* | 4.86E-05 | 1.15E-05 | 3.43E-05 | *RNH203* | 98 | *DDC1* | 88 |
| *RNH203* | 4.68E-05 | 6.79E-06 | 1.31E-05 | *MLH1* | 96 | *FUS2* | 88 |
| *YLR235C* | 4.52E-05 | 2.57E-06 | 6.11E-07 | *NUP170* | 96 | *HST3* | 88 |
| *TOF1* | 4.39E-05 | 1.40E-05 | 9.45E-04 | *PMS1* | 96 | *KIP1* | 88 |
| *YAP1* | 4.22E-05 | 5.04E-06 | 8.67E-06 | *ALE1* | 94 | *MFT1* | 88 |
| *RNH202* | 3.96E-05 | 1.38E-05 | 1.96E-03 | *APN1* | 94 | *MNT2* | 88 |
| *RNH201* | 3.86E-05 | 6.08E-06 | 1.91E-06 | *NFI1* | 94 | *MRPL51* | 88 |
| *SGS1* | 3.75E-05 | 1.42E-05 | 2.25E-03 | *YGR117C* | 94 | *NIT3* | 88 |
| *YDL162C* | 3.34E-05 | 9.73E-06 | 1.38E-03 | *YML020W* | 94 | *PCL10* | 88 |
| *PMS1* | 3.33E-05 | 1.28E-05 | 3.46E-03 | *YMR166C* | 94 | *PET123* | 88 |
| *HYR1* | 3.16E-05 | 1.74E-05 | 1.85E-02 | *YOR072W* | 94 | *PHM8* | 88 |
| *MGS1* | 3.10E-05 | 3.83E-06 | 6.14E-05 | *RPL23a* | 94 | *REC114* | 88 |
| *MSH2* | 3.09E-05 | 1.34E-06 | 1.55E-06 | *DIA2* | 92 | *RGS2* | 88 |
| *DST1* | 3.07E-05 | 6.56E-06 | 1.15E-04 | *EFT1* | 92 | *SCO1* | 88 |
| *YER188W* | 2.99E-05 | 1.27E-05 | 9.90E-03 | *MDM1* | 92 | *SPR1* | 88 |
| *CSM3* | 2.64E-05 | 3.65E-06 | 2.78E-04 | *MSN4* | 92 | *TOM5* | 88 |
| *HTA2* | 2.60E-05 | 6.24E-06 | 1.87E-03 | *PNS1* | 92 | *ULS1* | 88 |
| *RAD4* | 2.35E-05 | 2.46E-06 | 1.73E-03 | *RMI1* | 92 | *YDL009C* | 88 |
| *MSH6* | 2.34E-05 | 1.02E-05 | 1.68E-02 | *RRT14* | 92 | *YEL020C* | 88 |
| *RAD6* | 2.22E-05 | 7.25E-06 | 7.23E-03 | *SAC3* | 92 | *YGL042C* | 88 |
| *RRM3* | 2.16E-05 | 8.30E-06 | 1.54E-02 | *YDR230W* | 92 | *YJL017W* | 88 |
| *CHL4* | 2.14E-05 | 5.86E-06 | 9.36E-03 | *YLR235C* | 92 | *YJR018W* | 88 |
| *THP2* | 1.94E-05 | 2.95E-06 | 2.52E-03 | *YNL122C* | 92 | *YJR124C* | 88 |
| *DFG16* | 1.80E-05 | 4.48E-06 | 2.44E-02 | *YTA7* | 92 | *YKL091C* | 88 |
| *ABZ2* | 1.66E-05 | 3.93E-06 | 4.25E-02 | *FSH1* | 90 | *YKL162C* | 88 |
|  |  |  |  | *GET3* | 90 | *YNL179C* | 88 |
|  |  |  |  | *KGD2* | 90 | *YOR309C* | 88 |
|  |  |  |  | *MID2* | 90 | *YOR333C* | 88 |
|  |  |  |  | *POL32* | 90 |  |  |

<sup>a</sup> Recombination rate from Table S2
<sup>b</sup> p-values from one-sided Student's *t*-test

### Statistical analysis

Statistical analysis was performed in Excel or R (https://cran.r-project.org/).

### Data availability

Strains are available upon request. A file containing supplemental tables is available at FigShare. Table S1 lists all the strains used in this study. Table S2 contains the fluctuation test data from the patch screen. Table S3 contains the raw high-throughput replica pinning screen data. Table S4 contains the filtered pinning screen data. Table S5 contains the fluctuation test data from the pinning screen. Table S6 contains the GO term enrichment data.

## RESULTS

### A genetic screen for elevated spontaneous direct-repeat recombination

The *leu2* direct-repeat recombination assay (Smith and Rothstein, 1999) can detect both intra-chromosomal and sister chromatid recombination events (Figure 1A). Two nonfunctional *leu2* heteroalleles are separated by a 5.3 kb region containing the *URA3* marker. Reconstitution of a functional *LEU2* allele can occur either via sister chromatid recombination (gene conversion), which maintains the *URA3* marker, or via intra-chromosomal SSA, with the concomitant deletion of the sequence between the direct repeats and subsequent loss of the *URA3* marker (Symington et al., 2014). Both recombination events can be selected on media lacking leucine. We used this assay to systematically screen the yeast knockout (YKO) collection for genes whose deletion results in hyper-recombination between direct repeats (Figure 1B). We introduced the *leu2* direct-repeat recombination reporter into the YKO collection via synthetic genetic array (SGA) technology (Kuzmin et al., 2016). Each of the ∼4500 obtained strains was then patched on non-selective plates and replica-plated to plates lacking leucine to detect spontaneous recombination events as papillae on the replica-plated patches (Figure 1C). We included a wild-type control and a hyper-recombinant *elg1Δ* control (Bellaoui et al., 2003; Ben-Aroya et al., 2003) on every plate for reference. The recombination rates for 195 putative hyper-rec mutants identified by replica-plating (Table S2) were measured by a fluctuation test. Strains with a recombination rate greater than 2×10^-5^ (approximately twofold of the wild-type rate; 38 strains) were assayed in triplicate (or more). Thirty-three gene deletion mutant strains with a statistically supported increase in direct-repeat recombination rate relative to the wild-type control were identified (Figure 1D, Table S2, Table 1). The genes identified showed a high degree of enrichment for GO terms reflecting roles in DNA replication and repair (Figure 1E).

### A high-throughput screen for altered spontaneous direct-repeat recombination

We recently developed a high-throughput replica-pinning method to detect low-frequency events, and validated the scheme in a genome-scale mutation frequency screen (Novarina et al., 2020). To complement the data obtained with the classical screening approach, and to test our new methodology independently of the replicative aging context in which it was developed, we applied it to detect changes in spontaneous direct-repeat recombination (Figure 2A). We again introduced the *leu2* direct-repeat recombination reporter (Figure 1A) into the YKO collection. The collection was then amplified by parallel high-throughput replica-pinning to yield 48 colonies per gene deletion strain. After one day of growth, all colonies were replica-pinned (twice, in series) to media lacking leucine to select for recombination events. Recombination frequencies (a proxy for the spontaneous recombination rate) were calculated for each strain of the collection (Figure 2B, Table S3, Table S4). As a reference, recombination frequencies for the wild type (46%) and for a recombination-deficient *rad54Δ* strain (21%) obtained in a pilot replica-pinning experiment of 3000 colonies are indicated. In the screen itself, where 48 colonies were assessed, the wild type (*his3Δ::kanMX*) had a recombination frequency of 56%. Notably, a group of strains from the YKO collection carry an additional mutation in the mismatch repair gene *MSH3* (Lehner et al., 2007). Given the elevated spontaneous recombination rates of several mismatch repair-deficient strains (Figure 1D), we suspected that these *msh3* strains would display increased recombination frequencies, independently of the identity of the intended gene deletion. Indeed, the distribution of recombination frequencies for *msh3* strains (median: 74%) is shifted toward higher values compared to the overall distribution of the YKO collection (median: 60%) (Figure 2B). The 73 *msh3* strains were excluded from further analysis.

**Figure 2.**
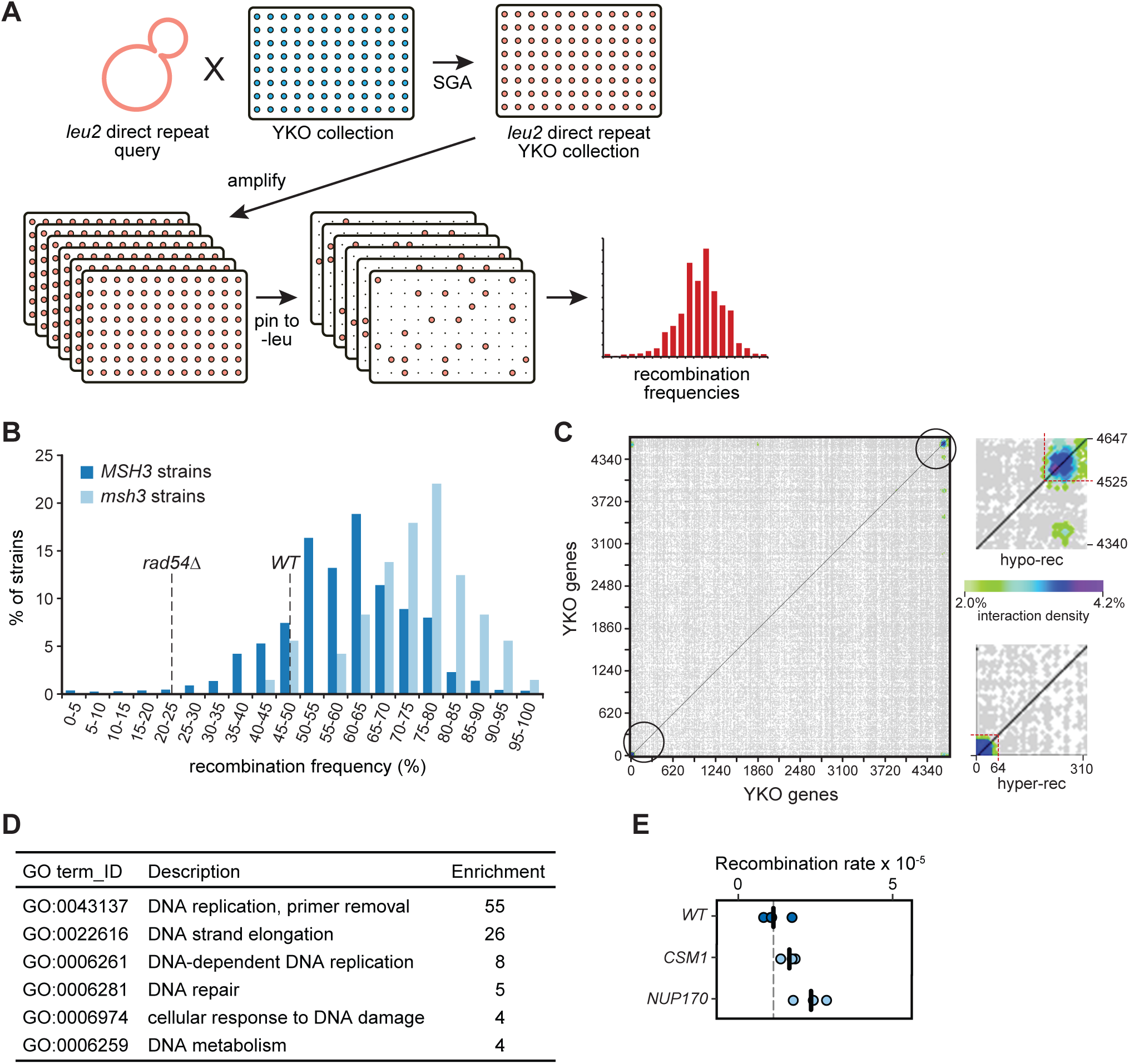
A high-throughput replica-pinning screen for genes controlling direct-repeat recombination. (**A**) Schematic representation of the screen based on high-throughput replica-pinning. The *leu2* direct-repeat recombination cassette was introduced into the yeast deletion collection as in Figure 1B. The resulting strains were amplified by parallel high-throughput replica pinning and subsequently replica-pinned to media lacking leucine to select for recombination events. Recombination frequencies were calculated for each strain of the YKO collection. (**B**) Recombination frequency distribution for the YKO collection (*MSH3* strains) and for the *msh3* strains in the collection. Recombination frequencies for a wild-type and for a recombination-defective *rad54Δ* strain derived from a pilot experiment are indicated by the dashed lines. (**C**) Interaction densities determined by CLIK analysis are plotted as a two-dimensional heatmap. The cutoffs established by CLIK analysis for hyper-recombination (hyper-rec) and recombination-defective (hypo-rec) genes are shown in the insets. (**D**) The statistically supported GO terms enriched in the hits from the pinning assay screen are shown, with the enrichment for each term. (**E**) Recombination rates from fluctuation tests of *csm1Δ* and *nup170Δ* are plotted. Each data point is from an independent fluctuation test, with n=3 for each strain. The vertical bars indicate the mean recombination rate for each strain and the wild-type data from Figure 1D are plotted for comparison.

To explore the overall quality of the high-throughput replica-pinning screen and to determine a cutoff in an unbiased manner, we performed Cutoff Linked to Interaction Knowledge (CLIK) analysis (Dittmar et al., 2013). The CLIK algorithm identified an enrichment of highly interacting genes at the top and at the bottom of our gene list (ranked according to recombination frequency), confirming the overall high quality of our screen, and indicating that we were able to detect both hyper- and hypo-recombinogenic mutants (Figure 2C). The cutoff indicated by CLIK corresponds to a recombination frequency of 87% for the hyper-recombination strains (75 genes; Table 1), and of 33% for the recombination-deficient strains (122 genes; Table 2).

**Table 2.**
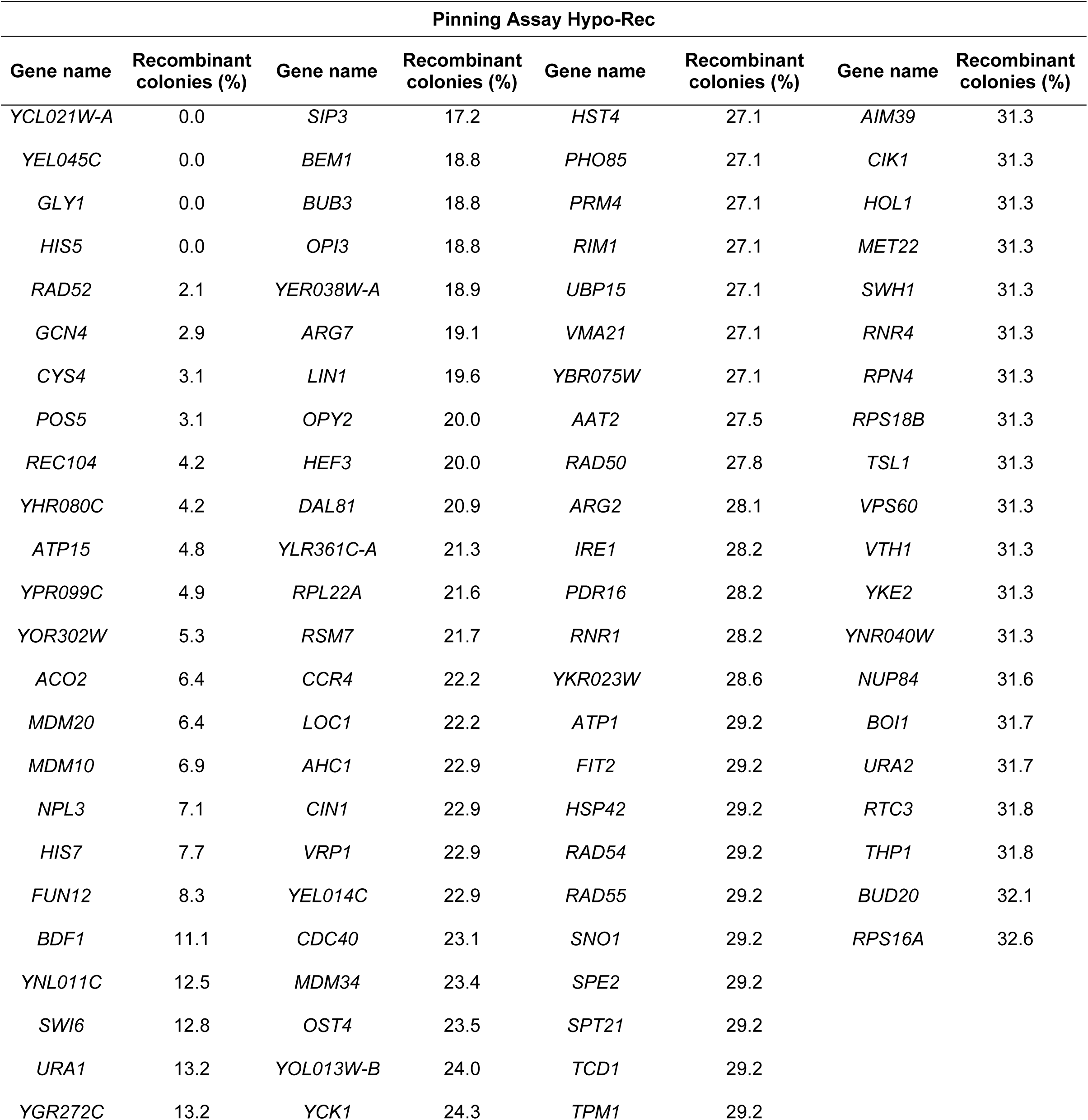
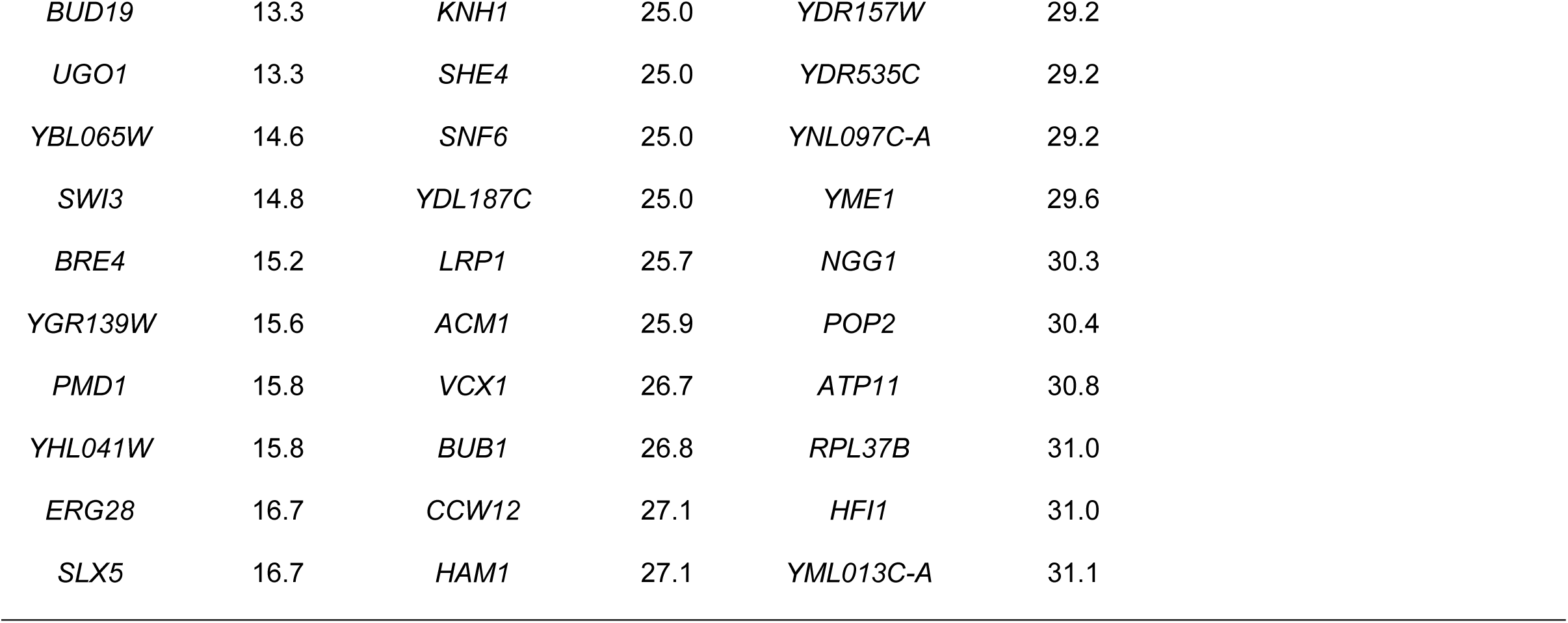
Hypo-recombination genes from the pinning assay screen.

| Pinning Assay Hypo-Rec |  |  |  |  |  |  |  |
| --- | --- | --- | --- | --- | --- | --- | --- |
| Gene name | Recombinant colonies (%) | Gene name | Recombinant colonies (%) | Gene name | Recombinant colonies (%) | Gene name | Recombinant colonies (%) |
| *YCL021W-A* | 0.0 | *SIP3* | 17.2 | *HST4* | 27.1 | *AIM39* | 31.3 |
| *YEL045C* | 0.0 | *BEM1* | 18.8 | *PHO85* | 27.1 | *CIK1* | 31.3 |
| *GLY1* | 0.0 | *BUB3* | 18.8 | *PRM4* | 27.1 | *HOL1* | 31.3 |
| *HIS5* | 0.0 | *OPI3* | 18.8 | *RIM1* | 27.1 | *MET22* | 31.3 |
| *RAD52* | 2.1 | *YER038W-A* | 18.9 | *UBP15* | 27.1 | *SWH1* | 31.3 |
| *GCN4* | 2.9 | *ARG7* | 19.1 | *VMA21* | 27.1 | *RNR4* | 31.3 |
| *CYS4* | 3.1 | *LIN1* | 19.6 | *YBR075W* | 27.1 | *RPN4* | 31.3 |
| *POS5* | 3.1 | *OPY2* | 20.0 | *AAT2* | 27.5 | *RPS18B* | 31.3 |
| *REC104* | 4.2 | *HEF3* | 20.0 | *RAD50* | 27.8 | *TSL1* | 31.3 |
| *YHR080C* | 4.2 | *DAL81* | 20.9 | *ARG2* | 28.1 | *VPS60* | 31.3 |
| *ATP15* | 4.8 | *YLR361C-A* | 21.3 | *IRE1* | 28.2 | *VTH1* | 31.3 |
| *YPR099C* | 4.9 | *RPL22A* | 21.6 | *PDR16* | 28.2 | *YKE2* | 31.3 |
| *YOR302W* | 5.3 | *RSM7* | 21.7 | *RNR1* | 28.2 | *YNR040W* | 31.3 |
| *ACO2* | 6.4 | *CCR4* | 22.2 | *YKR023W* | 28.6 | *NUP84* | 31.6 |
| *MDM20* | 6.4 | *LOC1* | 22.2 | *ATP1* | 29.2 | *BOI1* | 31.7 |
| *MDM10* | 6.9 | *AHC1* | 22.9 | *FIT2* | 29.2 | *URA2* | 31.7 |
| *NPL3* | 7.1 | *CIN1* | 22.9 | *HSP42* | 29.2 | *RTC3* | 31.8 |
| *HIS7* | 7.7 | *VRP1* | 22.9 | *RAD54* | 29.2 | *THP1* | 31.8 |
| *FUN12* | 8.3 | *YEL014C* | 22.9 | *RAD55* | 29.2 | *BUD20* | 32.1 |
| *BDF1* | 11.1 | *CDC40* | 23.1 | *SNO1* | 29.2 | *RPS16A* | 32.6 |
| *YNL011C* | 12.5 | *MDM34* | 23.4 | *SPE2* | 29.2 |  |  |
| *SWI6* | 12.8 | *OST4* | 23.5 | *SPT21* | 29.2 |  |  |
| *URA1* | 13.2 | *YOL013W-B* | 24.0 | *TCD1* | 29.2 |  |  |
| *YGR272C* | 13.2 | *YCK1* | 24.3 | *TPM1* | 29.2 |  |  |

|  |  |  |  |  |  |
| --- | --- | --- | --- | --- | --- |
| *BUD19* | 13.3 | *KNH1* | 25.0 | *YDR157W* | 29.2 |
| *UGO1* | 13.3 | *SHE4* | 25.0 | *YDR535C* | 29.2 |
| *YBL065W* | 14.6 | *SNF6* | 25.0 | *YNL097C-A* | 29.2 |
| *SWI3* | 14.8 | *YDL187C* | 25.0 | *YME1* | 29.6 |
| *BRE4* | 15.2 | *LRP1* | 25.7 | *NGG1* | 30.3 |
| *YGR139W* | 15.6 | *ACM1* | 25.9 | *POP2* | 30.4 |
| *PMD1* | 15.8 | *VCX1* | 26.7 | *ATP11* | 30.8 |
| *YHL041W* | 15.8 | *BUB1* | 26.8 | *RPL37B* | 31.0 |
| *ERG28* | 16.7 | *CCW12* | 27.1 | *HFI1* | 31.0 |
| *SLX5* | 16.7 | *HAM1* | 27.1 | *YML013C-A* | 31.1 |

**Table 3.**
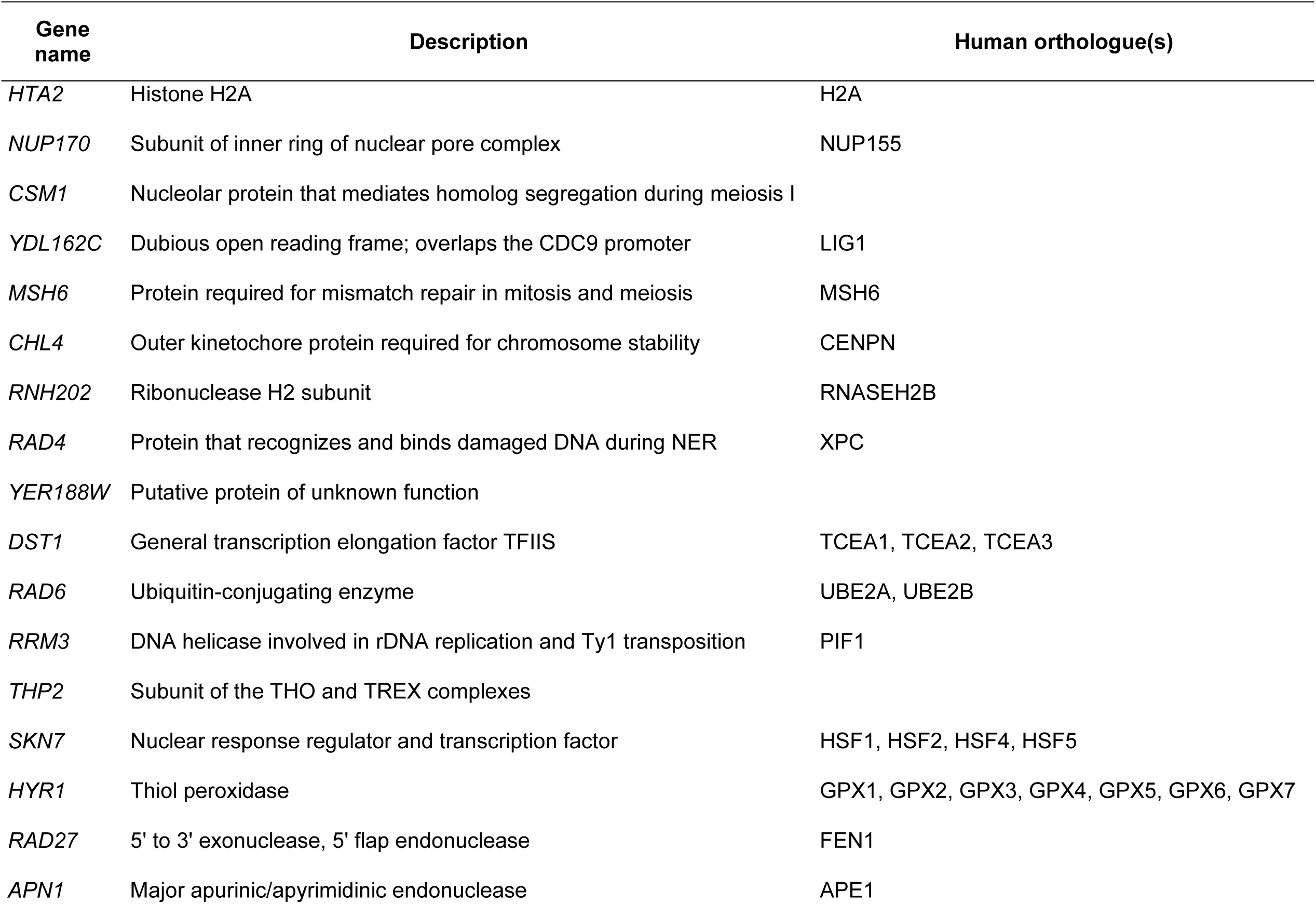
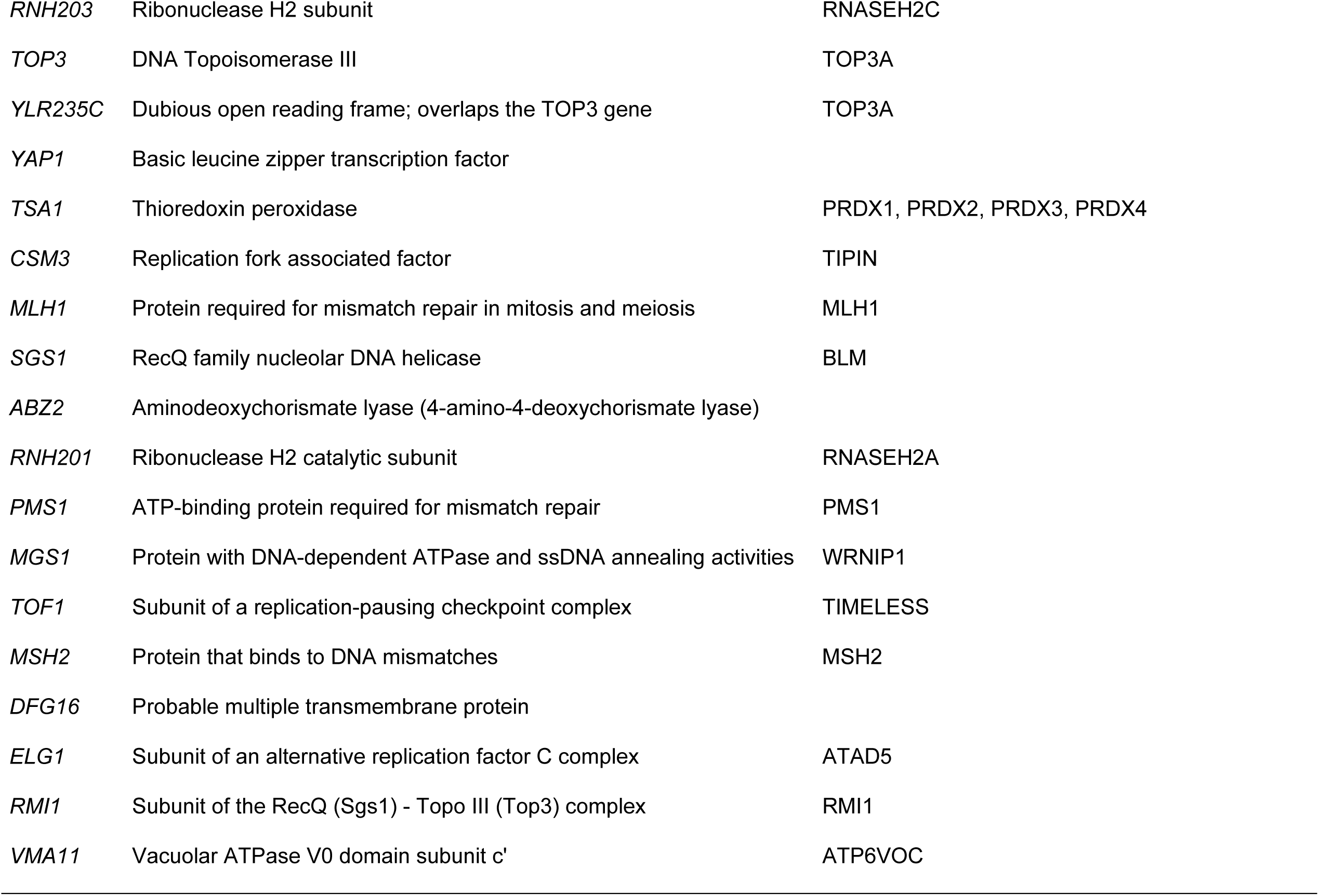
Validated hyper-recombination genes from the patch assay and pinning assay screens.

#### Hyper-recombination genes

We assessed the functions of the 75 hyper-recombination genes identified by our high-throughput screen (Figure 2D). As with the genes identified in the patch screen, the genes identified in the pinning screen were enriched for DNA replication and repair functions. Most importantly, at the very top of our hyper-recombination gene list (with 96% to 100% recombination), 11 out of 13 genes were identified in the patch screen and validated by fluctuation analysis (Table S2). We tested the two additional genes, *CSM1* and *NUP170*, by fluctuation analysis, and found that both had a statistically supported increase in recombination rate (Figure 2E and Table S5). Eighteen validated hyper-recombination genes from the patch screen were not identified in the pinning screen, and so are false negatives. Although we have not validated the weaker hits from the pinning screen (those with recombination frequencies between 87% and 96%), four genes in this range were validated as part of the patch screen (*APN1*, *RMI1*, *YLR235C*, and *RNH201*), 9 caused elevated levels of Rad52 foci when deleted (*APN1*, *NFI1*, *RMI1*, *POL32*, *RNH201*, *DDC1*, *HST3*, *MFT1*, and *YJR124C*) (Alvaro et al., 2007; Styles et al., 2016), and 3 are annotated as ‘mitotic recombination increased’ (*RMI1*, *DDC1*, and *HST3*; *Saccharomyces* Genome Database). Together these data suggest that additional bona fide hyper-recombination genes were identified in the pinning screen.

#### Hypo-recombination genes

By contrast to the replica-plating screen, the pinning screen detected mutants with reduced recombination frequency, with 122 genes identified (Table 2). The genes identified were functionally diverse, with no gene ontology (GO) processes enriched. Only 19 nonessential genes are annotated as having reduced recombination as either null or hypomorphic alleles in the *Saccharomyces* genome database (SGD; accessed January 11, 2020 via YeastMine). Of these, three genes (*RAD52*, *LRP1*, and *THP1*) were detected in the pinning screen. In addition, other members of the *RAD52* epistasis group important for effective homologous recombination (*RAD50*, *RAD54* and *RAD55*) displayed a recombination frequency lower than 33%, and *RAD51* was just above the cutoff (Table S3). Thus, our high-throughput replica-pinning approach detects mutants with very low recombination frequencies. More generally, this observation suggests that if the pinning procedure is properly calibrated, a high-throughput replica-pinning screen is able not only to detect mutants with increased rates of a specific low-frequency event (in this case direct-repeat recombination), but also mutants with reduced rates of the same low-frequency event.

#### Validated hyper-recombination genes identified in both screens

We compared the genes identified in the pinning screen with those identified in the patch screen, revealing 15 genes that were identified in both screens, a statistically supported enrichment (Figure 3A; hypergeometric p = 1.2×10^-21^). Combining the results of the two screens, we validated 35 genes whose deletion results in elevated spontaneous direct-repeat recombination (Table 3). Analysis of the group of 35 hyper-rec genes revealed 68 pairwise protein-protein interactions (Figure 3B), with many cases where several (if not all) members of the particular protein complex were identified. We found that 29 of the hyper-rec genes had at least one human orthologue (Table 3), indicating a high degree of conservation across the 35 validated genes. To assess the functional properties of the 35 gene hyper-rec set, we applied spatial analysis of functional enrichment (SAFE) (Baryshnikova, 2016) to determine if any regions of the functional genetic interaction similarity yeast cell map (Costanzo et al., 2016) are over-represented for the hyper-rec gene set (Figure 3C). We found a statistically supported over-representation of the hyper-rec genes in the DNA replication and repair neighbourhood of the genetic interaction cell map, highlighting the importance of accurate DNA synthesis in suppressing recombination. Finally, we compared the validated hyper-rec genes to relevant functional genomic instability datasets (*Saccharomyces* Genome Database annotation, (Alvaro et al., 2007; Hendry et al., 2015; Stirling et al., 2011; Styles et al., 2016); Figure 3D). Eight of our hyper-rec genes (*HTA2*, *MSH6*, *YER188W*, *ABZ2*, *PMS1*, *MSH2*, *DFG16*, and *VMA11*) were not identified in these datasets, indicating that our screens identified uncharacterized recombination genes. *HTA2*, *MSH6*, *PMS1*, *MSH2* have recombination phenotypes reported (see Discussion). Thus, we identify four genes without a characterized role in preventing recombination: *YER188W*, *ABZ2*, *DFG16*, and *VMA11*.

**Figure 3.**
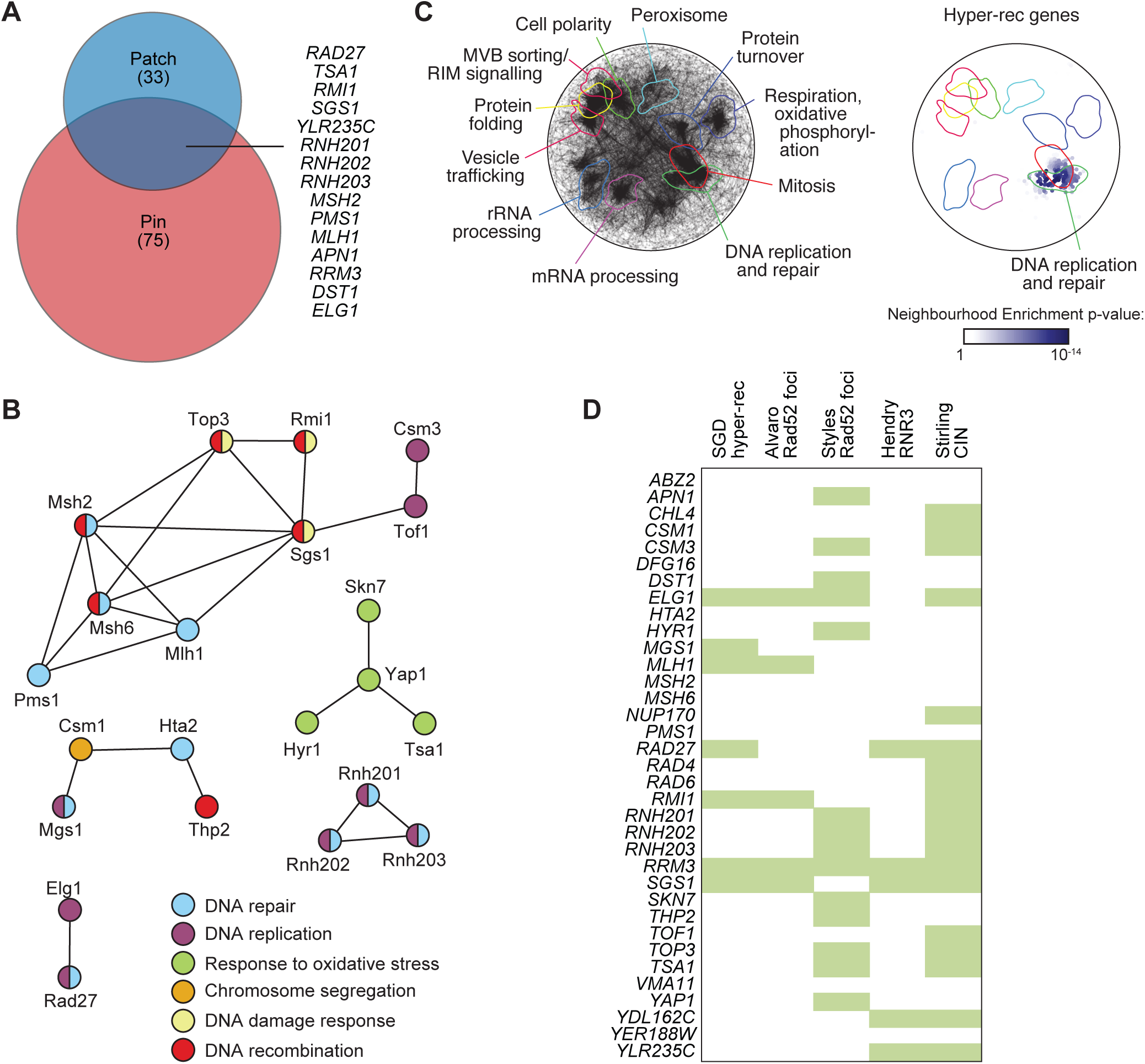
Functional analysis of validated hyper-rec genes. **(A)** The overlap of the hyper-rec genes for the two screens is plotted as a Venn diagram. The 15 genes identified in both screens are indicated. **(B)** A protein-protein interaction network for the proteins encoded by the 35 validated hyper-rec genes is shown. Nodes represent the proteins, and are colored to indicate function. Edges indicate a physical interaction as annotated in the GeneMania database. **(C)** Spatial analysis of functional enrichment. On the left, the yeast genetic interaction similarity network is annotated with GO biological process terms to identify major functional domains (Costanzo et al. 2016). 11 of the 17 domains are labeled and delineated by coloured outlines. On the right, the network is annotated with the 35 validated hyper-rec genes. The overlay indicates the functional domains annotated on the left. Only nodes with statistically supported enrichments (SAFE score > 0.08, p < 0.05) are coloured. (**D**) The 35 validated hyper-rec genes are compared with existing *Saccharomyces* Genome Database annotations and genome instability datasets that measured Rad52 focus formation (Alvaro et al., 2007; Styles et al., 2016), *RNR3* induction (Hendry et al., 2015), or chromosome instability (CIN; (Stirling et al., 2011)). A green bar indicates that the gene has the given annotation or was detected in the indicated screen.

To infer gene function for the four genes lacking a characterized role in suppressing recombination, we again applied SAFE analysis (Baryshnikova, 2016) to annotate the functional genetic interaction similarity yeast cell map (Costanzo et al., 2016) to identify any regions that are enriched for genetic interactions with each of the four genes (Figure 4). Of particular interest, the mitochondrial functional neighbourhood is enriched for negative genetic interactions with *YER188W* (Figure 4), suggesting that deletion of *YER188W* confers sensitivity to mitochondrial dysfunction. Analysis of *DFG16* revealed enrichments for positive interactions in the RIM signaling neighbourhood, which is expected (Barwell et al., 2005), but also for negative interactions in the DNA replication region of the map (Figure 4), indicating that *DFG16* is important for fitness when DNA replication is compromised. Analysis of *VMA11* revealed enrichment in the vesicle trafficking neighbourhood, typical of vacuolar ATPase subunit genes, and analysis of *ABZ2* revealed little (Figure 4). We conclude that functional analysis suggests mechanisms by which loss of *YER188W* (oxidative stress) or *DFG16* (genome integrity) results in hyper-recombination.

**Figure 4.**
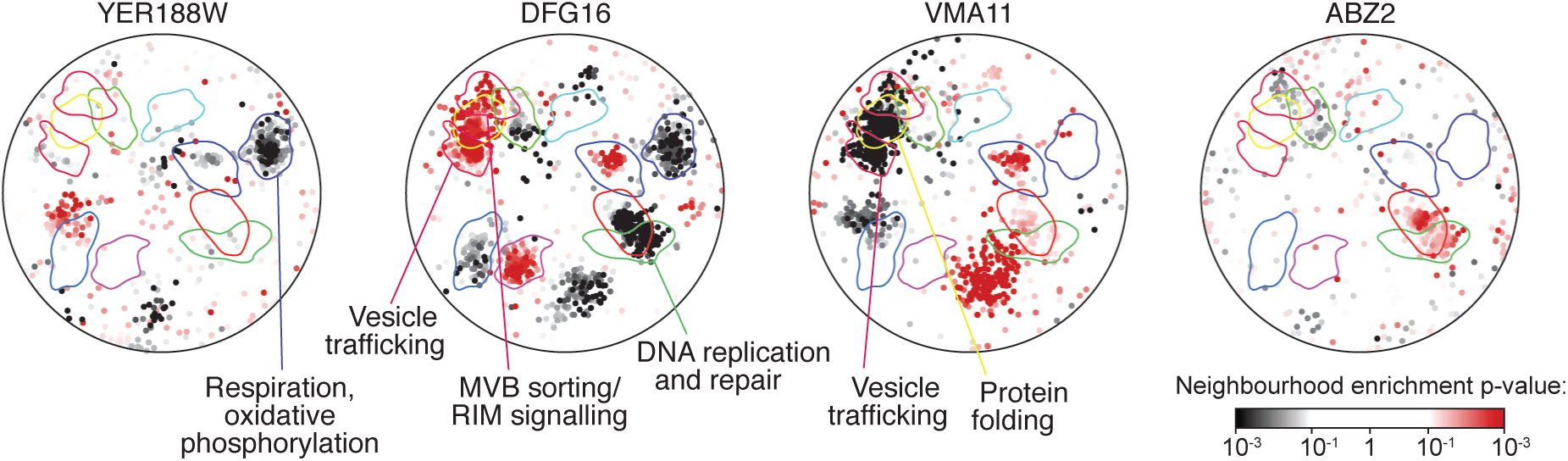
Spatial analysis of functional enrichment for four hyper-rec genes. The genetic interactions of each of the indicated genes was tested for enrichments in the functional neighbourhoods of the yeast genetic interaction similarity network. The overlay indicates a subset of functional domains as annotated on Figure 3C. Nodes with statistically supported enrichments (Neighbourhood enrichment p < 0.05) are coloured, black for negative genetic interactions and red for positive genetic interactions.

## DISCUSSION

Here we briefly discuss the functions of the genes and complexes identified in the screens and subsequently validated by fluctuation analysis.

### Mismatch repair

*MLH1*, *MSH2*, *MSH6* and *PMS1* are evolutionary conserved genes involved in mismatch repair (MMR), a pathway that detects and corrects nucleotide mismatches in double-strand DNA (Spies and Fishel, 2015). An anti-recombinogenic role for these four MMR genes in yeast has been previously described: specifically, MMR proteins are important to prevent homeologous recombination and SSA between slightly divergent sequences, via mismatch recognition and heteroduplex rejection (Datta et al., 1996; Nicholson et al., 2000; Spies and Fishel, 2015; Sugawara et al., 2004). The role for MMR in preventing homeologous recombination is conserved also in mammalian cells (de Wind et al., 1995; Elliott and Jasin, 2001; Spies and Fishel, 2015). It is worth noting that the presence of sequence differences between the two *leu2* alleles in the *leu2* direct-repeat assay is essential to genetically detect recombination events. Therefore, it is reasonable that this assay should detect genes involved in suppressing homeologous recombination.

### Sgs1-Top3-Rmi1 complex

The evolutionary conserved helicase-topoisomerase complex Sgs1-Top3-Rmi1 is involved in DSB resection and in dissolution of recombination intermediates (Symington et al., 2014). Consistent with previous observations (Chang et al., 2005), our screen identified all three members of the complex, together with *YLR235C*, a dubious ORF that overlaps the *TOP3* gene. The Sgs1-Top3-Rmi1 complex dissolves double Holliday junction structures to prevent crossover formation (Cejka et al., 2010). The same role has been reported for BLM helicase, the human Sgs1 homolog mutated in the genome stability disorder Bloom syndrome (Wu et al., 2006; Yang et al., 2010). Furthermore, several genetic studies indicate that the anti-recombinogenic activity of Sgs1-Top3-Rmi1 cooperates with MMR proteins in heteroduplex rejection to prevent homeologous recombination (Chakraborty et al., 2016; Goldfarb and Alani, 2005; Myung et al., 2001; Spell and Jinks-Robertson, 2004; Sugawara et al., 2004).

### MGS1

In our screen we also identified *MGS1*, the homolog of the WRN-interacting protein WRNIP1. Mgs1 displays DNA-dependent ATPase and DNA strand annealing activities. Deletion of *MGS1* causes hyper-recombination, including elevated direct-repeat recombination (Hishida et al., 2001). It seems that Mgs1 promotes faithful DNA replication by regulating Pol*δ*, and promoting replication fork restart after stalling (Branzei et al., 2002; Saugar et al., 2012). The absence of Mgs1 could result in increased replication fork collapse, leading to the formation of recombinogenic DSBs (Branzei et al., 2002). Similar roles have been suggested for WRNIP1 in mammalian cells (Leuzzi et al., 2016; Tsurimoto et al., 2005).

### RNase H2 complex

*RNH201* encodes the evolutionary conserved catalytic subunit of RNase H2, while the two non-catalytic subunits are encoded by *RNH202* and *RNH203* genes. This enzyme cleaves the RNA moiety in RNA-DNA hybrids originating from Okazaki fragments, co-transcriptional R-loops, and ribonucleotide incorporation by replicative polymerases (Cerritelli and Crouch, 2009). Deletion of any of the three subunits in yeast inactivates the whole complex. Human RNase H2 genes are mutated in Aicardi-Goutières syndrome, a severe neurological disorder (Crow et al., 2006). Inactivation of yeast RNase H2 causes elevated LOH, ectopic recombination and direct-repeat recombination (Conover et al., 2015; Potenski et al., 2014), mostly dependent on Top1 activity. What is the recombinogenic intermediate accumulated in the absence of RNase H2? It has been suggested that Top1-dependent cleavage at the ribonucleotide site creates a nick that can be further converted into a recombinogenic DSB (Potenski et al., 2014). Recent genetic studies indicate that, while in the case of LOH events hyper-recombination is caused by Top1-dependent processing of single ribonucleotides incorporated by leading strand polymerases and/or by accumulation of recombinogenic R-loops (Conover et al., 2015; Cornelio et al., 2017; Keskin et al., 2014; O’Connell et al., 2015), elevated direct-repeat recombination results instead from Top1-dependent cleavage of stretches of ribonucleotides, resulting from defective R-loop removal or Okazaki fragment processing in the absence of RNase H2 (Epshtein et al., 2016). In line with this model, we also detected elevated direct-repeat recombination rate in the absence of the Thp2 member of the THO complex, which functions at the interface between transcription and mRNA export to prevent R-loop accumulation (Chavez et al., 2000; Huertas and Aguilera, 2003), *DST1*, which encodes a transcription elongation factor and is anti-recombinogenic (Owiti et al., 2017), and the flap endonuclease encoded by *RAD27*, which is involved in Okazaki fragment processing (Balakrishnan and Bambara, 2013) (Table 3).

Furthermore, deletion of the dubious ORF *YDL162C*, also identified in our screen, likely affects the expression level of neighbouring *CDC9*, an essential gene encoding DNA Ligase I, involved in Okazaki fragment processing and ligation after ribonucleotide removal from DNA. Together, available data suggest that different modes leading to accumulation of RNA-DNA hybrids or unprocessed Okazaki fragments result in hyper-recombination.

### Fork protection complex

Tof1 and Csm3 (Timeless and Tipin in human cells) form the fork protection complex (FPC), involved in stabilization of replication forks, maintenance of sister chromatid cohesion and DNA replication checkpoint signaling (Bando et al., 2009; Chou and Elledge, 2006; Katou et al., 2003; Leman et al., 2010; Mayer et al., 2004; Mohanty et al., 2006; Noguchi et al., 2004, 2003; Xu et al., 2004). Recently, Tof1 and Csm3 were implicated in restricting fork rotation genome-wide during replication; they perform this role independently of their interacting partner Mrc1, which we did not identify in our screen (Schalbetter et al., 2015). In the absence of Tof1 or Csm3, excessive fork rotation can cause spontaneous DNA damage, in the form of recombinogenic ssDNA and DSBs (Chou and Elledge, 2006; Schalbetter et al., 2015; Sommariva et al., 2005; Urtishak et al., 2009). Indeed, depletion of Tof1 and Csm3 orthologues results in accumulation of recombination intermediates in fission yeast and mouse cells (Noguchi et al., 2004, 2003; Sommariva et al., 2005; Urtishak et al., 2009).

### RRM3

The *RRM3* gene, encoding a 5′ to 3′ DNA helicase, was initially identified because its absence causes hyper-recombination between endogenous tandem-repeated sequences (such as the rDNA locus and the *CUP1* genes) (Keil and McWilliams, 1993). The Rrm3 helicase travels with the replication fork and facilitates replication through genomic sites containing protein-DNA complexes that, in its absence, cause replication fork stalling and breakage. Such Rrm3-dependent sites include the rDNA, telomeres, tRNA genes, inactive replication origins, centromeres, and the silent mating-type loci (Azvolinsky et al., 2006; Ivessa et al., 2003, 2000; Schmidt and Kolodner, 2004; Torres et al., 2004). Intriguingly, a tRNA gene is located about 350 bp upstream the chromosomal location of the *leu2* direct-repeat recombination marker. Increased replication fork pausing in the absence of Rrm3 could cause recombinogenic DSBs, explaining the elevated direct-repeat recombination we observe in the *rrm3Δ* strain.

### Oxidative stress response genes

*YAP1* and *SKN7* encode two transcription factors important for the activation of the cellular response to oxidative stress (Morano et al., 2012). The glutathione peroxidase encoded by *HYR1* has a major role in activating Yap1 in response to oxidative stress (Delaunay et al., 2002). *TSA1* is a Yap1 and Skn7 target and encodes a peroxiredoxin that scavenges endogenous hydrogen peroxide (Wong et al., 2004). Deletion of *TSA1* causes hyper-recombination between inverted repeats (Huang and Kolodner, 2005), and oxidative stress response genes (including *TSA1*, *SKN7* and *YAP1*) are synthetic sick or lethal with HR mutants (Pan et al., 2006; Yi et al., 2016). A likely explanation for the elevated direct-repeat recombination we measured in strains defective for the oxidative stress response, therefore, is that oxidative DNA damage generates replication blocking lesions and/or replication-associated DSBs, both of which are processed by the HR pathway (Huang and Kolodner, 2005). An alternative explanation could be that extensive oxidative DNA damage results in the saturation of the mismatch-binding step of MMR, compromising MMR-dependent heteroduplex rejection, resulting in increased homeologous recombination (Hum and Jinks-Robertson, 2018; Spies and Fishel, 2015).

### Other DNA Repair genes

*APN1* encodes the main apurinic/apyrimidinic (AP) endonuclease involved in yeast base excision repair (BER). Removal of endogenous alkylating damage can generate abasic sites, which are mostly processed by Apn1 (Boiteux and Guillet, 2004; Popoff et al., 1990; Xiao and Samson, 1993). In the absence of *APN1*, abasic sites accumulate, which can hamper DNA replication. The recombination pathway is involved in the repair and/or bypass of these abasic sites, as suggested by the genetic interactions between the BER and the HR pathways (Boiteux and Guillet, 2004; Swanson et al., 1999; Vance and Wilson, 2001). The *APN1* gene is adjacent to *RAD27*, and therefore it is also possible that the hyper-recombination phenotype of *apn1Δ* is due to a “neighbouring-gene effect” on *RAD27*, as was reported in the case of telomere length alteration (Ben-Shitrit et al., 2012).

*HTA2*, which encodes one copy of histone H2A, is of course important for appropriate nucleosome assembly. Reducing histone levels by deleting one H3-H4 gene pair or by partial depletion of H4 increases recombination (Clemente-Ruiz and Prado, 2009; Liang et al., 2012; Prado and Aguilera, 2005), and it is likely that reducing *HTA2* gene dosage also does so. Since histone depletion results in diverse chromatin defects, the exact mechanisms by which recombination is induced are elusive.

*RAD4* encodes a key factor of nucleotide excision repair (NER), and is involved in direct recognition and binding of DNA damage (Prakash and Prakash, 2000), while *RAD6* is a key gene controlling the post replication repair (PRR) DNA damage tolerance pathway (Ulrich, 2005). Genetic studies suggest that BER, NER, PRR and HR can redundantly process spontaneous DNA lesions, and inactivation of one pathway shifts the burden on the others. This mechanism could explain why deletion of *RAD4* or *RAD6* causes a modest increase in spontaneous direct-repeat recombination (Swanson et al., 1999).

*CSM1* encodes a nucleolar protein that serves as a kinetochore organizer to promote chromosome segregation in meiosis, and is involved in localization and silencing of rDNA and telomeres in mitotic cells (Poon and Mekhail, 2011). Interestingly, Csm1 is important to inhibit homologous recombination at the rDNA locus and other repeated sequences (Burrack et al., 2013; Huang et al., 2006; Mekhail et al., 2008). The nuclear pore complex has an intimate connection to recombination, in that some DSBs move to and are likely repaired at the NPC (Freudenreich and Su, 2016). The NPC gene *NUP170* has not been directly implicated in DSB repair, but is important for chromosome segregation (Kerscher et al., 2001).

### The unknowns (YER188W, ABZ2, DFG16, and VMA11)

Unexpectedly, the top hyper-rec gene identified in our screen is *VMA11*, which encodes a subunit of the evolutionarily conserved vacuolar H^+^-ATPase (V-ATPase), important for vacuole acidification and cellular pH regulation (Hirata et al., 1997; Kane, 2006; Umemoto et al., 1991). *VMA11* involvement in genome maintenance is suggested by the sensitivity of a *vma11Δ* strain to several genotoxic agents, namely doxorubicin, ionizing radiation, cisplatin and oxidative stress (Thorpe et al., 2004; Xia et al., 2007). V-ATPase defects in yeast result in endogenous oxidative stress and defective Fe/S cluster biogenesis as a consequence of mitochondrial depolarization (Hughes and Gottschling, 2012; Milgrom et al., 2007; Veatch et al., 2009). Of note, several DNA replication and repair factors are Fe/S cluster proteins (Veatch et al., 2009; Zhang, 2014). Therefore, the hyper-recombination phenotype of *vma11Δ* could be due to increased spontaneous DNA damage, caused by elevated endogenous oxidative stress and/or by defective DNA replication and repair as a consequence of compromised Fe/S cluster biogenesis. However, *VMA11* was not detected in screens for increased Rad52 foci (Alvaro et al., 2007; Styles et al., 2016), or in a screen for increased DNA damage checkpoint activation (Hendry et al., 2015), suggesting that spontaneous DNA damage might not accumulate to high levels in *vma11Δ*.

*ABZ2* encodes an enzyme involved in folate biosynthesis (Botet et al., 2007). Folate deficiency and the resulting compromise of nucleotide synthesis could promote recombination, although yeast culture media are rich in folate, and the *ABZ2* genetic interaction profile reveals no similarity to nucleotide biosynthesis genes (Usaj et al., 2017). *DFG16* encodes a predicted transmembrane protein involved in pH sensing (Barwell et al., 2005). Interestingly, SAFE analysis indicates a role for *DFG16* in DNA replication and/or DNA repair, in addition to the expected role in pH signaling. There is currently little insight into the function of *YER188W*. SAFE analysis indicates a possible role in mitochondrial function, however a protein product of *YER188W* has not been detected to date in either mass spectrometry or GFP fusion protein analyses (Breker et al., 2014; Ho et al., 2018; Huh et al., 2003).

## ACKNOWLEDGMENTS

We thank Anastasia Baryshnikova for advice and assistance with the SAFE analysis. This work was supported by grants from the Netherlands Organisation for Scientific Research (Vidi grant 864.12.002 to MC) and the Canadian Institutes of Health Research (MOP-79368 and FDN-159913 to GWB).

